# Flagella methylation promotes bacterial adhesion and host cell invasion

**DOI:** 10.1101/774588

**Authors:** Julia A. Horstmann, Michele Lunelli, Hélène Cazzola, Johannes Heidemann, Caroline Kühne, Pascal Steffen, Sandra Szefs, Claire Rossi, Ravi K. Lokareddy, Chu Wang, Kelly T. Hughes, Charlotte Uetrecht, Hartmut Schlüter, Guntram A. Grassl, Theresia E.B. Stradal, Yannick Rossez, Michael Kolbe, Marc Erhardt

## Abstract

The flagellum is the motility device of many bacteria and the long external filament is made of several thousand copies of a single protein, flagellin. While posttranslational modifications of flagellin are common among bacterial pathogens, the role of lysine methylation remained unknown. Here, we show that both flagellins of *Salmonella enterica*, FliC and FljB, are methylated at surface-exposed lysine residues. A *Salmonella* mutant deficient in flagellin methylation was outcompeted for gut colonization in a gastroenteritis mouse model. In support, methylation of flagellin promoted invasion of epithelial cells *in vitro*. Lysine methylation increased the surface hydrophobicity of flagellin and enhanced flagella-dependent adhesion of *Salmonella* to phosphatidylcholine vesicles and epithelial cells. In summary, posttranslational flagellin methylation constitutes a novel mechanism how flagellated bacteria facilitate adhesion to hydrophobic host cell surfaces and thereby contributes to efficient gut colonization and successful infection of the host.

## Introduction

The Gram-negative enteropathogen *Salmonella enterica* uses a variety of strategies to successfully enter and replicate within a host. In this respect, bacterial motility enables the directed movement of the bacteria towards nutrients or the target site of infection. A rotary nanomachine, the flagellum, mediates motility of many bacteria, including *Salmonella enterica*^1^. Flagella also play a central role in other infection processes, involving biofilm formation, immune system modulation and adhesion^2–4^.

The eukaryotic plasma membrane plays an important role in the interaction of flagellated bacteria with host cells during the early stages of infection^5^. The flagella of *S. enterica, Escherichia coli* and *Pseudomonas aeruginosa* can function as adhesion molecules^6–8^ mediating the contact to various lipidic plasma membrane components, including cholesterol, phospholipids, sulfolipids and the gangliosides GM1 and aGM1^9–12^.

Structurally, the flagellum consists out of three main parts: the basal body embedded within the inner and outer membranes of the bacterium, a flexible linking structure - the hook, and the long, external filament, which functions as the propeller of the motility device^13^. The filament is formed by more than 20,000 subunits of a single protein, flagellin. Many *S. enterica* serovars express either of two distinct flagellins, FliC or FljB, in a process called flagellar phase variation^14,15^. FliC-expressing bacteria display a distinct motility behavior on host cell surfaces and a competitive advantage in colonization of the intestinal epithelia compared to FljB-expressing bacteria^16^. However, while the structure of FliC has been determined previously^17^, the structure of FljB remained unknown.

The many thousand surface-exposed flagellin molecules are a prime target of the host’s immune system. Accordingly, many flagellated bacteria have evolved mechanisms to prevent flagellin recognition, *e.g*. by posttranslational modifications of flagellin. Flagellin glycosylation is relatively common among *Enterobacteriaceae*^18^, in *Campylobacter, Aeromonas* and *Pseudomonas* species^19–21^ and plays a critical role in adhesion, biofilm formation or mimicry of host cell surface glycans^22,23^. *S. enterica* does not posttranslationally glycosylate its flagellins. However, *ε*-N-methylation at lysine residues of flagellin via the methylase FliB has been reported^24–26^. Although flagellin methylation was first reported in 1959^25^, the physiological role of the methylation remained elusive. Previous studies indicated that the absence of FliB had no significant effect on swimming and swarming motility suggesting that flagellin methylation might be required for virulence of *Salmonella^27,28^*.

In the present study, we analyzed the role of flagellin methylation for motility and virulence of *S. enterica in vivo* and *in vitro*. Our results demonstrate that *S. enterica* exploits methylated flagella to adhere to hydrophobic host cell surfaces. Thus, the posttranslational methylation of flagellin plays an important role for invasion of host cells, and accordingly, productive colonization of the host’s epithelium.

## Results

Previous studies suggested that the flagellins of *S. enterica* are posttranslationally methylated, however, the identity of the methylated lysine residues remained largely unknown^25,26,29^. We performed mass spectrometry analyses with high sequence coverage of both flagellins FliC and FljB isolated from *S. enterica* genetically locked in expression of FliC (*fliC*^ON^) or FljB (*fljB*^ON^), respectively, and isogenic mutants of the methylase FliB (Δ*fliB*) (Supplementary Fig. S1). In order to map the identified *ε*-N-methyl-lysine residues to the structure of both flagellins, we determined the crystal structure of FljB (Supplementary Fig. S2, Supplementary Text S1). The tertiary structure of FljB resembles, similar to FliC, a boomerang-shape with one arm formed by the D1 domain and the other formed by D2 and D3^17^. However, in FljB compared with FliC the variable D3 domain is rotated about 90° around the axis defined by the D2-D3 arm, resulting in the widening angle of about 20° between the two boomerang’s arms (Fig. 1a, Supplementary Fig. S2). Interestingly, the methylated lysine residues are primarily located in the surface-exposed D2 and D3 domains of both flagellins (Fig. 1a, Supplementary Fig. S2d). In FliC, 16 out of 28 lysine residues and in FljB, 19 out of 30 lysine residues were methylated. We note that the methylation of 15 lysines of FliC and 18 lysines of FljB was dependent on the presence of FliB, while only one lysine in FljB and one in FliC was methylated in the absence of FliB (Δ*fliB*). 10 of the identified lysines were methylated in both FliC and FljB flagellins, while 6 and 9 methylation sites were unique to FliC and FljB, respectively (Supplementary Fig. S2d). Interestingly, conserved lysines were methylated in a FliB-dependent manner in both flagellins except for Lys^396^ in FljB that is not modified in the corresponding Lys^385^ in FliC.

**Fig. 1:**
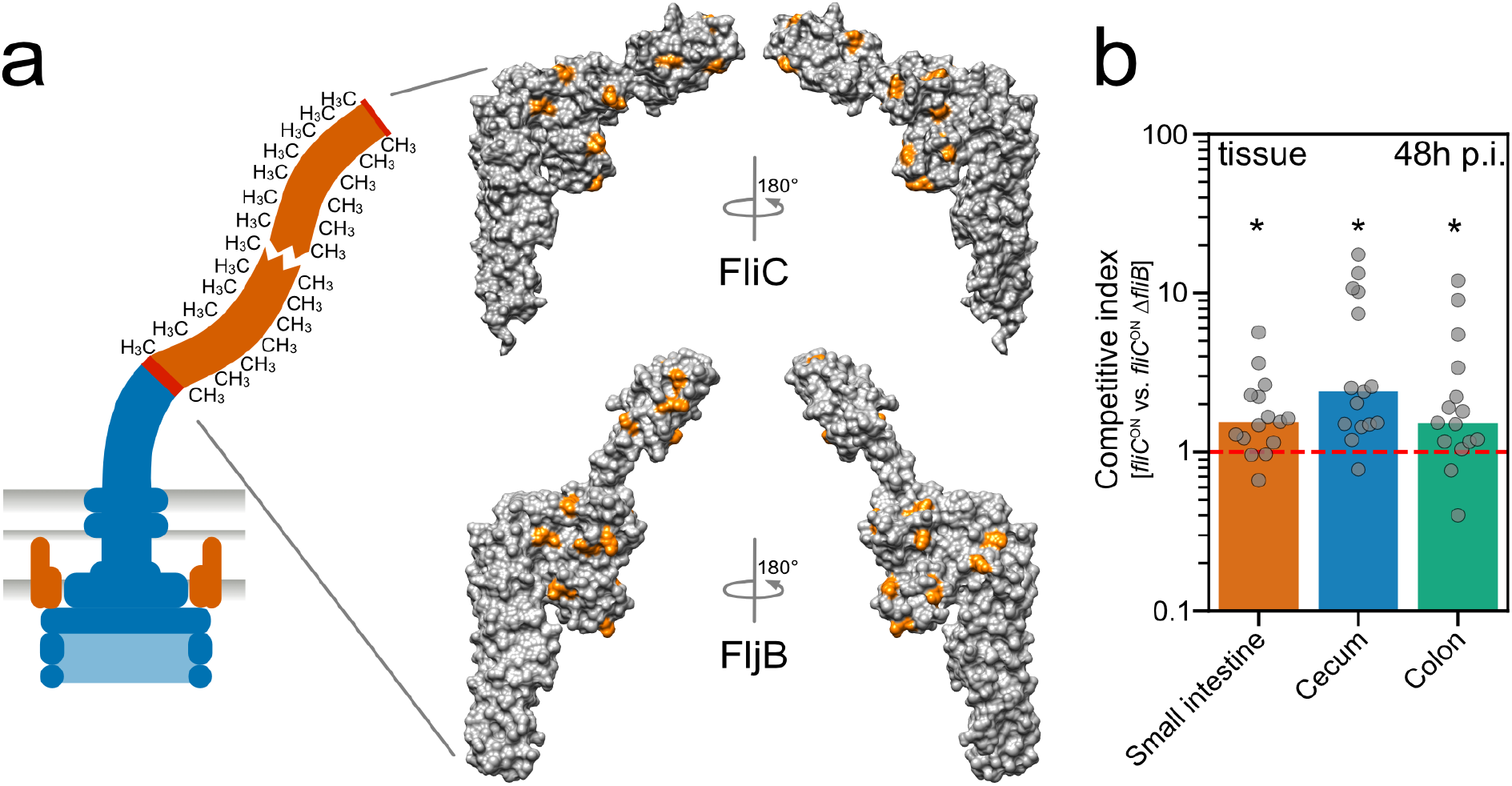
Surface-exposed methylation of flagellin contributes to efficient colonization of the murine intestine. (a) Schematic of a methylated flagellar filament and surface representation of the structure of FliC (top) and FljB (bottom). FliB-dependent methylation sites are highlighted in orange. (b) Streptomycin pre-treated C57BL/6 mice were infected with 10^7^ CFU of the FliC-expressing WT (*fliC*^ON^) and isogenic Δ*fliB* mutant, each harboring a different antibiotic resistant cassette. The organ burden (small intestine, colon and cecum lumen and tissue, respectively) was determined two days post-infection and used to calculate the competitive indices (CI). Each mouse is shown as an individual data point and significances were analyzed by the Wilcoxon Signed Rank test. The bar graph represents the median of the data and asterisks indicate a significantly different phenotype to the value 1 (* = *p*<0.05).

We next aligned the amino acid sequences of FljB and FliC up- and downstream of the identified ε-N-methyl-lysine residues (± 6 residues, Supplementary Fig. S3). Although no clear consensus sequences could be determined, we found prevalence of small (Ala, Gly, Thr, Val, Ser) and negatively charged (Asp) residues around the methylated lysines. Interestingly, a scan of the local sequences that surround methylated lysines using ScanProsite^30^ matched the profile of the bacterial Ig-like domain 1 (Big-1) for 11 and 10 FliB-dependent modifications in FljB and FliC, respectively, although with low confidence level (Supplementary Table S2). We note that the Big-1 domain is present in adhesion proteins of the intimin/invasin family, which are crucial in bacterial pathogenicity mediating host-cell invasion or adherence^31–33^.

Based on the weak homology of the methylation sites to the Big-1 domain and the absence of a motility phenotype in non-methylated flagellin mutants (Supplementary Fig. S4, Supplementary Text S2), we hypothesized that flagellin methylation might play a role in *Salmonella* virulence. We thus co-infected streptomycin-pre-treated mice^34^ with the wildtype (WT) and an isogenic Δ*fliB* mutant (Fig. 1b). Organ burden analysis two days post-infection revealed that the Δ*fliB* strain was significantly outcompeted by the WT in the gastroenteritis mouse model, especially in the cecal tissue (Fig. 1b, competitive indices >1), suggesting that methylated flagella play an important role for efficient colonization of the intestinal epithelium.

We next tested if invasion of epithelial cells *in vitro* was also dependent on flagellin methylation (Fig. 2a). We first infected murine MODE-K epithelial cells with the WT and *S. enterica* strains deficient in the methylase FliB and determined the number of intracellular bacteria. Invasion was reduced about 50% for the Δ*fliB* mutant strain independently of the flagellin type (Fig. 2b, top). We also observed a similar invasion defect for the Δ*fliB* mutant when we forced contact of the bacteria with the epithelial cells using centrifugation (Fig. 2b, bottom), suggesting that the invasion defect of the Δ*fliB* mutant did not depend on active bacterial motility.

**Fig. 2:**
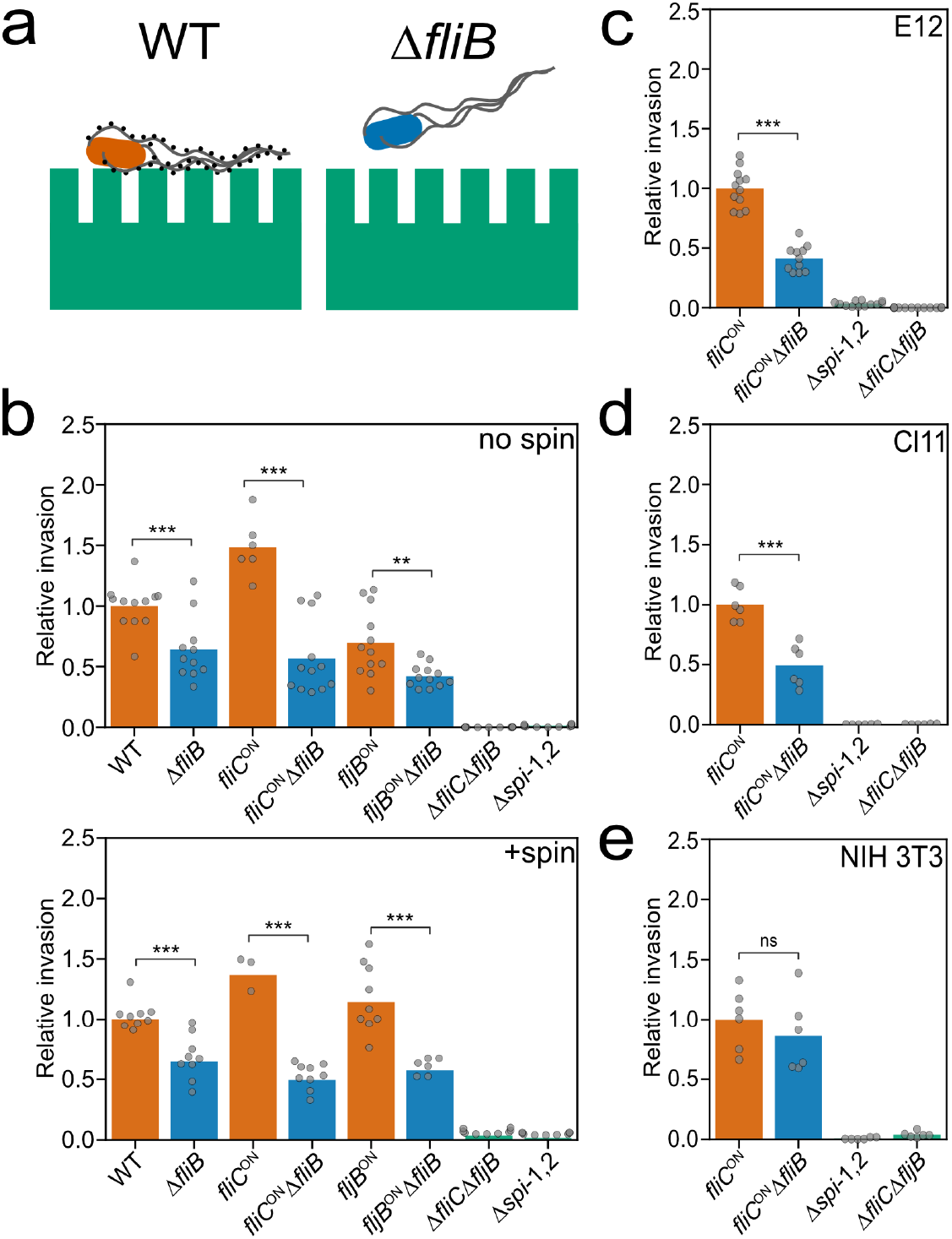
Flagella methylation facilitates eukaryotic cell invasion. (a) Schematic illustration of productive adhesion and invasion of eukaryotic epithelial cells dependent on methylated flagella. (b) Invasion of MODE-K murine epithelial cells depends on methylated flagella. Reported are relative invasion rates of MODE-K epithelial cells for various flagellin methylation mutants without (top: no spin) or with forced contact of the bacteria by centrifugation (bottom: +spin). (c-e) Relative invasion rates of different eukaryotic host cell types. The human epithelial cell line E12, the murine epithelial cell line Cl11, and the murine fibroblast cell line NIH 3T3 were infected with *Salmonella* flagella methylation mutants as described above. The bar graphs represent the mean of the reported relative invasion rate data normalized to the inoculum. Replicates are shown as individual data points and statistical significances were determined by the Student’s *t* test (** = P<0.01; *** = P<0.001; ns = not significant).

We further analyzed if the methylation-dependent invasion phenotype was eukaryotic cell-line specific (Fig. 2c-e, Supplementary Fig. S5). The human epithelial cell line E12^35^ and murine intestinal epithelial cell line Cl11 mimic the native intestinal environment *in vitro*. Similar to the observed invasion rate of MODE-K cells, a Δ*fliB* mutant strain displayed a two-fold decreased invasion rate of the human and murine epithelial cell lines compared with the WT. Similarly, in murine epithelial-like RenCa cells, the invasion rate of a Δ*fliB* mutant was decreased. Invasion into the murine fibroblast cell lines NIH 3T3 and CT26, however, was independent of flagellin methylation, suggesting that the observed invasion phenotype is cell type-specific for epithelial-like cells.

We next confirmed that the observed invasion phenotype was due to the lack of *fliB* by complementing expression of *fliB* from an inducible Ptet promoter at its native chromosomal locus. Addition of anhydrotetracycline (AnTc) induced *fliB* expression comparable to levels of the WT and restored invasion of MODE-K epithelial cells (Supplementary Fig. S6). We further tested if the observed invasion defect was dependent on the assembly of the methylated flagellar filament. A hook deletion mutant *(ΔflgE)* does not express flagellin, whereas a mutant of the hook-filament junction proteins *(ΔflgKL*) expresses and secretes flagellin, but does not assemble the flagellar filament. The methylase FliB is expressed in both Δ*flgE* and Δ*flgKL* mutant backgrounds^27^. We observed in neither the Δ*flgE* nor the Δ*flgKL* mutant a difference in MODE-K epithelial cell invasion in the presence or absence of FliB, suggesting that methylated flagellin must assemble into a functional flagellar filament in order to facilitate epithelial cell invasion (Supplementary Fig. S7).

Our results presented above demonstrate that the presence of an assembled, methylated flagellar filament, but not the ability to move *per se*, contributes to the observed defect of *Salmonella* to invade epithelial cells. We thus hypothesized that adhesion to epithelial cells was facilitated through methylated flagella. In particular, we reasoned that the addition of hydrophobic methyl groups to surface-exposed lysine residues (Fig. 1) would affect the hydrophobicity of the flagellar filament. Consistently, the surface hydrophobicity So of purified FliC and FljB flagella was significantly reduced in the absence of lysine methylation (Fig. 3a+b, Supplementary Fig. S8). These results suggested that methylated flagella might promote adhesion to hydrophobic molecules present on the surface of epithelial cells. We therefore investigated adhesion of *S. enterica* to MODE-K epithelial cells. In order to dissect flagella methylation-dependent adhesion from methylation-dependent invasion of the epithelial cells, we employed *Salmonella* mutants deleted for the *Salmonella* pathogenicity island-1 (spi-1), which renders the bacteria unable to invade epithelial cells in an injectisome-dependent manner. We found that adhesion of Δ*spi-1 Salmonella* mutants to MODE-K epithelial cells was reduced up to 50% for strains deficient in flagellin methylation (Fig. 3c). This result suggested that methylated flagella promote adhesion to the hydrophobic surface of epithelial cells or surface-exposed proteinaceous receptors or glycostructures.

**Fig. 3:**
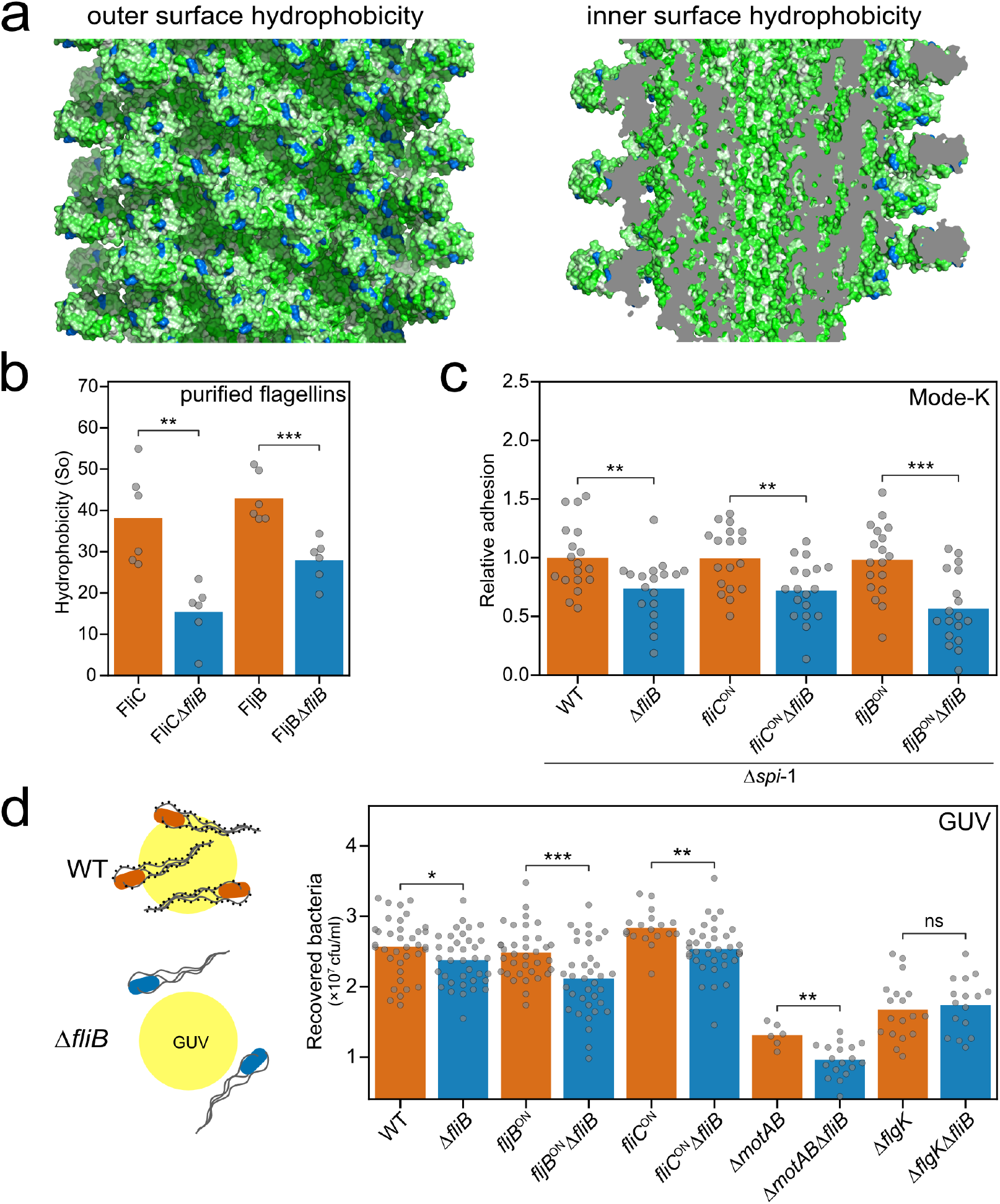
Flagella methylation mediates adhesion to hydrophobic surfaces. (a) Methylation increases hydrophobicity of the flagellar filament outer surface. Surface hydrophobicity distribution of the outer (left) and the inner surface (right) of the FliC flagellar filament^36^ according to the Eisenberg scale^37^ (from green to white indicates increasing hydrophobicity) with FliB-dependent methylation sites highlighted in blue. (b) Measured surface hydrophobicity (So) of methylated and non-methylated *(ΔfliB)* flagellins using PRODAN on purified flagellar filaments. (c) Adhesion of *S. enterica* to MODE-K epithelial cells is reduced in the absence of flagella methylation. Adhesion was monitored using *S. enterica* strains deleted for *spi-1* in order to prevent invasion of the eukaryotic host cells. (d) Adhesion of *S. enterica* to giant unilamellar vesicles (GUV) consisting of phosphatidylcholine from egg chicken is dependent on the presence of methylated flagella. Left: schematic illustration of the adhesion of *Salmonella* to GUVs dependent on methylated flagella. Right: Quantified adhesion of *Salmonella* mutants to GUVs. The bar graphs represent the mean of the reported data. Replicates are shown as individual data points and statistical significances were determined by the Student’s *t* test (* = P<0.05; ** = P<0.01; *** = P<0.001; ns = not significant).

However, we did not observe a significant flagella methylation-dependent effect on adhesion of *Salmonella* to various extracellular matrix proteins, nor to the oligosaccharide mannose, which has previously been shown to mediate adhesion of *Salmonella* and *E. coli* to eukaryotic cells using type I fimbriae^38–41^ (Supplementary Fig. S9). We next tested the possibility that the increased hydrophobicity of methylated flagella might promote adhesion of the bacteria to the hydrophobic plasma membrane of epithelial cells (Fig. 3d). We therefore analyzed the binding of *Salmonella* to giant unilamellar vesicles (GUV) consisting of phosphatidylcholine, the most abundant phospholipid in animal tissues. Interestingly, we observed a reduction in bacterial adhesion to the GUV for *S. enterica* strains deficient in flagellin methylation, but not for the non-flagellated Δ*flgK* mutants. In addition, non-motile, but flagellated bacteria *(ΔmotAB)* were less adherent, which supports previous observations that actively rotating flagella are important for the initial interaction with surfaces before biofilm formation^41^.

## Discussion

Flagella-dependent motility is crucial for *Salmonella* pathogenesis by enabling directed movement towards host epithelial cells. However, flagella not only play a role in bacterial motility, but also in colonization, adhesion, and biofilm formation^41–44^. In case of flagella-mediated adhesion to host cells, the primary interactions with the epithelial tissue occur *via* the external filament made of several thousand copies of flagellin and thus represents an excellent adhesion molecule.

Here, we describe a novel mechanism of flagella-dependent adhesion to surface-exposed hydrophobic molecules. This adhesion phenotype is dependent on methylation of surface-exposed lysine residues of flagellin by the methylase FliB. Flagellin methylation was first described in *Salmonella* in 1959^24–26^, however, the physiological relevance remained elusive. We demonstrate that FliB-mediated flagellin methylation is crucial for *Salmonella* pathogenesis in the mouse model and contributes significantly to adhesion and thus invasion of epithelial cells *in vitro*, but does neither affect swimming motility nor flagella assembly (Supplementary Text S3). Analysis of the surface hydrophobicity of purified flagella revealed that methylation of the filament subunits increases the hydrophobicity of the outer surface of the flagellar filament, while the lumen of the flagellar filament seems not to be affected (Fig. 3a+b, Supplementary Fig. S8). We note that the preferential methylation of surface-exposed lysine residues implicates a FliB methylation mechanism involving flagellin assemblies formed in the bacterial cytosol prior to secretion. Further, we found that a single flagellin molecule contains 16 or 19 surface-exposed methylation sites. Since a flagellar filament is made up of up to 20,000 flagellin copies, the methylation of flagellin subunits might substantially increase the overall hydrophobicity of the flagellum. Consistently, we found that adhesion to the surface of epithelial host cells and phospholipid vesicles was dependent on the flagella methylation status. In support, flagella have been recently implicated to mediate adhesion to abiotic surfaces through hydrophobic interactions^45,46^. We thus speculate that bacteria use flagella to explore the host cell surface as suggested previously^47^ and actively rotating flagella might be able to penetrate the lipid bilayer and interact with the fatty acids buried inside the plasma membrane. Increasing the surface hydrophobicity of the flagellar filament through methylation might improve those hydrophobic interactions for productive adhesion to eukaryotic host cells.

Flagellin Methylation Islands (FMI) and thus modifications of flagellins by methylation are common in *Enterobacteriaceae*^18^. In addition to *Salmonella*, many bacterial species including *Yersinia, Enterobacter, Franconibacter*, and *Pantoea* contain chromosomal FMI loci, which encode orthologues of FliB. In summary, FliB-dependent methylation of flagella might represent a general mechanism facilitating adhesion to hydrophobic host cell surfaces in a broad range of bacterial species.

## Methods

### Ethics statement

All animal experiments were performed according to guidelines of the German Law for Animal Protection and with permission of the local ethics committee and the local authority LAVES (Niedersächsisches Landesamt für Verbraucherschutz und Lebensmittelsicherheit) under permission number 33.19-42502-04-13/1191.

### Strains, media and bacterial growth

All bacterial strain used in this study are listed in Supplementary Table S3 and were derived from SL1344. Bacteria were grown in lysogeny broth (LB)^48^ at 37 °C and growth was measured by optical density at 600 nm. For transductional crosses the generalised transducing phage P22 *HT105/1 int-201* was used^49^. Gene deletions or replacements were produced as previously described^50^. All bacterial strains are available upon request.

### Cloning and purification of FljB for structural analysis

The truncated gene *fljB* encoding for the protein residues 55-462 was amplified from S. Typhimurium (SL1344) by standard PCR method and cloned into the expression vector pET28a(+) using the restriction sites NheI and XhoI to obtain N-terminal His-tagged protein. The mutation A190V was found by sequencing. Standard conditions were applied to express His-tagged FljB55-462 in BL21(DE3). The protein was purified from the soluble fraction using HisTrap HP and Superdex 75 columns (GE Healthcare) in 50 mM HEPES (pH 7.4), 150 mM NaCl.

### Crystallization and data collection

FljB_55-462_ was concentrated to 12-15 mg/mL and crystals were grown at 18 °C by hanging drop vapour diffusion against 0.1 M Tris (pH 8.5), 20% (w/v) PEG4000, 24% (v/v) isopropanol. Diffraction data were collected using crystals flash-frozen in crystallization buffer. Measurements were carried out at the beamLine BL14.1 at the Helmholtz-Zentrum Berlin synchrotron Bessy II, at the wavelength 0.918 Å and temperature 100 K, obtaining a data set at 2.00 Å resolution. Crystals belong to space group C2, with one FljB molecule in the asymmetric unit (solvent content 51.6%). Indexing, integration, merging and scaling were done using the program XDS^51^.

### Crystal structure determination

The structure was phased by molecular replacement, using the structure of the F41 fragment of FliC flagellin as search model (PDB ID 1IO1^17^). Cycles of manual building and refinement using Coot^52^ and CNS version 1.3^53^ led to the final structure, which includes residues 55-459 of FljB with the mutation A190V and the residue S54 present in the crystallised construct. 299 water molecules were also placed. Structural comparison between FljB and FliC has been done with the server PDBeFold v2.59^54^. Molecular structure figures were generated using UCSF Chimera 1.13.1^55^ and PyMol 2.2.3 (Schroedinger, LLC (2018). Alignment Fig. S2d was generated with the server ESPript (https://espript.ibcp.fr)^56^.

### Mass spectrometry

#### Sample preparation

FliC and FljB purified from the WT and a Δ*fliB* mutant were separated using SDS-PAGE. The corresponding bands were cut from the gel and each was cut into 1×1 mm pieces. An in-gel digestion was performed. For destaining the gel pieces, a 50 mM ammonium bicarbonate (AmBiCa) in 50% acetonitrile (ACN) solution was added and incubated 30 min at room temperature to dehydrate the gel. After removal of the supernatant, 50 mM AmBiCa was added and incubated for 30 min at room temperature to rehydrate the pieces. This step was repeated two times. Disulfide bonds were reduced using 10 mM DTT in 50 mM AmBiCa for 30 min at 56 °C. After cooling to room temperature and removal of the supernatant, the reduced cysteines were alkylated using 55 mM iodoacetamide in 50 mM AmBiCa for 30 min at room temperature in the dark. After removal of the supernatant the gel pieces were dried *in vacuo*. Ice cold 50 mM AmBiCa in 10% ACN containing 12.5 ng/*μ*L trypsin was added and digested overnight. Peptides were extracted by transferring the supernatant to a fresh collection tube and adding 50 mM AmBiCa in 10% ACN to the gel pieces and transferring the second supernatant into the same collection tube. Peptides were dried *in vacuo* and stored at −20 °C. Before measuring the peptides were reconstituted in 10 *μ*L 0.1% formic acid (FA) and 1 *μ*L was injected for measurement. All chemicals were purchased from Sigma-Aldrich. Trypsin was purchased from Promega.

#### Mass Spectrometry

Peptides were measured on a tandem mass spectrometer (Fusion, Thermo Fisher Scientific) coupled to a nano UPLC system (Ultimate 3000 RSLCnano, Thermo Fisher Scientific) with a nano-spray source. Peptides were trapped on a C18 reversed-phase trap column (2 cm x 75 *μ*m ID; Acclaim PepMap trap column packed with 3 *μ*m beads, Thermo Fisher Scientific) and separated on a 25 cm C18 reversed-phase analytical column (25 cm x 75 *μ*m ID, Acclaim PepMap, 3 *μ*m beads, Thermo Fisher Scientific). The column temperature was kept constant at 45 °C. Peptides were separated using a 2-step gradient starting with 3% buffer B (0.1% FA in ACN) and 97% buffer A (0.1% FA in H_2_O) with a steady increase to 28% buffer B over 20 min and a second increase to 35% over 5 min with a subsequent ramping to 90% buffer B for 10 min followed by a 20 min equilibration to 3% buffer B at a constant flow rate of 300 nL/min. Eluting peptides were injected directly into the mass spectrometer. Data were acquired in positive ion mode using data dependent acquisition (DDA) with a precursor ion scan resolution of 1. 2×10^5^ at 200 *m/z* in a range of 300-1500 *m/z* with an automatic gain control (AGC) target of 2×10^5^ and a maximum injection time of 50 ms. Peptides were selected for fragmentation using the “TopSpeed” method with a threshold of 5000 intensity and a dynamic exclusion time of 30 sec. Peptides were fragmented using higher-energy collision dissociation (HCD) in the C-Trap and fragment spectra were detected in the ion trap. Fragment spectra were recorded using the “Rapid” setting with a maximum injection time of 35 ms and an AGC target of 1×10^4^ with the first mass set at 110 *m/z*.

#### Data analysis

Data were analyzed using the ProteomeDiscoverer 2.0 (Thermo Fisher Scientific) software. Spectra were identified using the Sequest HT search engine with precursor mass tolerance set to 10 ppm and the fragment mass tolerance set to 0.5 Da. Carbamidomethylation on cysteine was set as fixed modification and oxidation on methionine, acetylation on protein N-terminus as well as mono-, di- and tri-methylation on lysine were set as variable modifications. Trypsin was set as enzyme and 3 missed cleavages were allowed with a minimum peptide length of 6 amino acids. Spectra were searched against a *Salmonella* Typhimurium FASTA database obtained from UniProt in June 2016 containing 1821 entries and a contaminant database containing 298 entries. Sequence coverage maps were created using PatterLab for proteomics 4.0^57^.

### Protein secretion assay

Protein secretion into the culture supernatant was analyzed as described before^16^. Samples were fractionated under denaturizing conditions on SDS-gels (200 OD units) and immunoblotting was performed using primary a-FliC/FljB and secondary a-rabbit antibodies.

### Motility assay and immunostaining of flagella

Swimming motility was analyzed in semi-solid agar plates as described before^58^. For immunostaining of flagella, logarithmically grown cells were fixed on a poly-L-lysine coated coverslip by 2% formaldehyde and 0.2% glutaraldehyde. Flagellin was stained using polyclonal α-FliC (rabbit, 1:1000 in 2% BSA/PBS) and secondary a-rabbit AlexaFluor488 (goat, 1:1000 in PBS). DNA was stained using DAPI (Sigma-Aldrich). Images were taken as described before^28,59^.

### Mouse infection studies

Intragastrical infection of seven weeks old C57BL/6 mice (Janvier) was performed as described in^16^. Briefly, mice were infected with 10^7^ CFU of two strains containing an antibiotic resistance cassette. Small intestine, cecum and colon were isolated 2 days post-infection and competitive indices (CI) were calculated.

### Invasion and adhesion assays

The murine epithelial cell lines MODE-K^60^ and Cl11, the murine epithelial-like cell line Renca (CRL-2947), the human epithelial cell line HT29-MTX-E12 (E12)^35^, and the mouse fibroblast cell lines NIH-3T3 (CRL-1658) and CT26 (CRL-2638) were used for invasion assays. The immortalization and characterization of the muGob (Cl11) cells will be described elsewhere (Truschel et al., in preparation). Briefly, murine intestinal organoids were plated and infected with different lentiviruses encoding the CI-SCREEN gene library^61^. After transduction, the clonal cell line muGob (Cl11) was established, which has integrated the following recombinant genes of the CI-SCREEN library: Id1, Id2, Id3, Myc, Fos, E7, Core, Rex (Zfp42). The muGob (Cl11) cell line was cultivated on fibronectin/collagen-coated (InSCREENeX GmbH, Germany) well plates in a humidified atmosphere with 5% CO2 at 37 °C in a defined muGob medium (InSCREENeX GmbH, Germany). 2.5×10^5^ cells/mL were seeded in 24-well plates. *Salmonella* strains were added for infection at a MOI of 10 for 1 h. External bacteria were killed by addition of 100 *μ*g/mL gentamycin for 1 h and cells were lysed with 1% Triton X-100. Serial dilutions of the lysate were plated to calculate the CFU/mL. All values were normalised to the control strain. To test adhesion to MODE-K cells, cells were seeded and infected with strains lacking *spi-1* to prevent injectisome-dependent invasion. After infection, the MODE-K cells were washed extensively and lysed as described above.

### RNA isolation and quantitative real-time PCR

Strains were grown under agitating growth conditions in LB medium and total RNA isolation was performed using the RNeasy Mini kit (Qiagen). For removal of genomic DNA, RNA was treated with DNase using the TURBO DNA-free kit (Ambion). Reverse transcription and quantitative real-time PCRs (qRT-PCR) were performed using the SensiFast SYBR No-ROX One Step kit (Bioline) in a Rotor-Gene Q Lightcycler (Qiagen). Relative changes in mRNA levels were analyzed according to Pfaffl^62^ and as described before^28^.

### ECM adhesion assays

For ECM protein adhesion assays, a 96-well plate pre-coated with a variety of ECM proteins was used (EMD Millipore; Collagen I, II, IV, Fibronectin, Laminin, Tenascin, Vitronectin). Wells were rehydrated according to the user’s manual and 5×10^7^ cells/mL were added. After incubation for 1h at 37 °C, wells were washed extensively and 1% Triton X-100 was added. Colony forming units (CFU)/mL were calculated after plating of serial dilutions and normalised to the inoculum and BSA control.

### Mannose binding assay

Binding to mannose was determined as described before^63^ with minor modifications. A black 96-well plate was coated with BSA or mannose-BSA (20 *μ*g/mL in 50 mM bicarbonate buffer pH 9.5) for 2 h at 37 °C, followed by blocking with BSA (10 mg/mL) for 1 h at 37°C. Adjusted bacterial cultures (OD_600_ 0.6) harboring the constitutive fluorescent plasmid pFU228-P_*gapdh*_-mCherry were added to the wells to facilitate binding. After 1 h incubation at 37°C, wells were washed with 1x PBS and fluorescence was measured with a Tecan plate reader (excitation 560 nm; emission 630 nm). Fluorescence relative to the binding to BSA was calculated from three technical replicates and the type-I fimbriae-inducible strain P_tet_-*fimA-F* served as positive control.

### Flagellar filament isolation

Flagellar filaments were isolated similar as described^64^. Briefly, bacterial cultures were grown in LB media at 37 °C for 16 h in an orbital shaker incubator (Infors HT) at 80 rpm. Cells were harvest by centrifugation at 2,000 x g for 20 min. Cell pellets were resuspended in TBS buffer pH 7.4 at 4 °C. The flagella were sheared off with a magnetic stirrer at 500 rpm for 1 h, followed by centrifugation at 4,000 x g for 30 min. Supernatants were collected and ammonium sulfate was slowly added while stirring to achieve two-thirds saturation. After overnight incubation, the flagella were harvest by centrifugation at 15,000 x g for 20 min and pellet was re-suspended in TBS buffer at pH 7.4. The quality of the purified flagella was checked by SDS-PAGE and transmission electron microscopy of negatively stained samples (microscope Talos L120C, Thermo Fisher Scientific).

### Hydrophobicity determination

Protein surface hydrophobicity was measured according to a modification of the method of Kato and Nakai^65^ using PRODAN^66^. A stock solution of 1mM PRODAN (prepared in DMSO) was used, 8 *μ*L was added to successive samples containing 1 mL of diluted flagella in 20 mM HEPES (pH 7.4), 150 mM NaCl. After homogenization by pipetting, the samples were incubated 10 min in the dark and the relative fluorescence intensity was measured. All fluorescence measurements were made with a Cary Eclipse (Varian now Agilent) spectrofluorometer. Excitation and emission wavelengths were 365 nm and 465 nm, the slit widths were 5 and 5 nm. For standardization, BSA was used. Surface hydrophobicity (So) values were determined using at least duplicate analyses. Five measures per sample repeated three times were performed and the mean was used.

### Bacterial adhesion to liposomes

Giant unilamellar vesicles (GUV) were prepared according to the polyvinyl alcohol (PVA)-assisted swelling method^67^. Gold-coated glass slides were obtained by thermal evaporation under vacuum (Evaporator Edwards model Auto 306, 0.01 nm·s^-1^, 2-3 × 10^6^ mbar). A gold layer of 10 ± 1 nm was deposited on top of a chromium adhesion layer of 1 ± 0.5 nm. Prior to GUV formation in HEPES buffered saline solution (HEPES 20 mM pH 7.4, NaCl 150 mM), 1,2-distearoyl-sn-glycero-3-phosphoethanolamine-N-[PDP(polyethylene glycol)-2000] (DSPE-PEG-PDP) (Sigma-Aldrich) was mixed with L-α-phosphatidylcholine from egg chicken (Sigma-Aldrich) at a 3 % mass ratio that allows a direct covalent coupling of GUV onto gold surfaces. For the bacterial adhesion assay, a 5 *μ*g/mL GUV solution was deposited onto a gold-coated glass substrate and incubated one hour for immobilization. Then, the surface was gently rinsed with buffer to remove non-immobilized liposomes. Subsequently bacterial culture at 10^8^ CFU/mL resuspended in HEPES buffer was carefully deposit on the surface and incubated for one hour. Non-adherent bacteria were eliminated by buffer washes. Finally, the adherent bacteria were detached by pipetting several times directly onto the immobilized liposomes with PBS pH 7.4. The collected samples were serially diluted and plated on LB agar for viable bacterial counts. Averages and standard deviations were calculated from six independent experiments.

### Data availability

The data that support the findings of this study are available from the corresponding authors upon request. The coordinates of the flagellin FljB have been deposited in the RCSB PDB under accession numbers 6RGV.

## Acknowledgements

We thank Heidi Landmesser, Nadine Körner, Henri Galez, Laurine Lemaire and Pauline Adjadj for expert technical assistance, Juana de Diego and members of the Erhardt and Kolbe labs for useful discussions and for critical comments on the manuscript and Keichi Namba for providing providing the atomic model of the FliC flagellar filament. We thank HZB for the allocation of synchrotron radiation beamtime and Uwe Müller for the support at the beamline BL14.1, Petra Dersch for kindly providing plasmid pFU228, Michael Hensel for providing P_tet_-*fimA-F* mutant strains and Tobias May (InSCREENeX GmbH) for help in tissue culture and providing the epithelial-like cell line Cl11.

## Author contributions

J.A.H., M.L., M.K. and M.E. conceived of the project, designed the study, and wrote the paper; J.A.H., M.L, H.C., J.H. and C.K. performed the experiments; J.A.H., M.L, H.C., J.H., C.K., C.U., G.A.G, Y.R., M.K. and M.E. analyzed the data; P.S., S.S., C.R., R.K.L., C.W. and K.T.H. contributed to experiments and performed strain construction; C.U., H.S., G.A.G., T.E.B.S., Y.R., M.K. and M.E. contributed funding and resources.

