## Supplementary Materials for "Flagella methylation promotes bacterial adhesion and host cell invasion"

Supplementary Text S1 – S3

Supplementary Figures S1 – S8

Supplementary Tables S1 – S3

### Supplementary Text:

#### Supplementary Text S1: Determining the structure of the flagellin FljB

Large parts of the different flagellin variants are conserved. In particular, the N- and C-termini of flagellin that form the D0 and D1 domains are highly conserved and necessary for formation of a coiled-coil structure and flagellin polymerization [1–3]. The D2 and D3 domains that form the horizontal arm of the molecule are surface-exposed in the assembled filament structure. These domains are highly variable and immunogenic, leading to recognition by host receptors and triggering of innate and adaptive immune responses, and the structural differences may also contribute to the distinct motility behavior [4–6]. However, while the structure of the F41 fragment of FliC, missing the D0 domain, has been determined previously [7], the structure of FljB remained unknown.

Accordingly, we determined the crystal structure of FljB from *S. enterica* (Supplementary Table S1). Despite several attempts, we were not able to crystallize full-length $FljB_{1\text{-}506}$ most likely due to the spontaneous polymerization caused by the D0 domain. We therefore crystallized a FljB fragment lacking the D0 domain based on the crystal structure of the F41 fragment of *Salmonella* FliC [7]. Deletion of the N- and C-terminal regions (comprising residues 1-54 and 463-506, respectively) of the D0 domain yielded crystals diffracting X-rays to 2.0 Å resolution. The crystallographic asymmetric unit contains a single copy of FljB comprising wildtype residues 55-459 in a continuous polypeptide chain and an additional serine residue in position 54 belonging to the tag inserted for affinity purification.

FljB can be divided into three domains named D1, D2 and D3 according to the structure (Fig. S2a). The topology of the FljB domains is conserved compared with FliC (Fig. S2b) [7]. Both the N- and C-termini contribute to the domain D1 with the polypeptide chain segments 55-177 and 414-459. D2 is also formed by two discontinuous segments of the polypeptide chain (residues 178-190 and 288-413) followed by domain D3 that is most distant from the FljB termini (residues 191-287) (Fig. S2b+d).

The tertiary structure of FljB resembles, similar to FliC, a boomerang with one arm formed by the D1 domain and the other formed by D2 and D3 (Fig. S2a). The major difference between the structures of the two flagellins is the rotation of the D3 domain of about 90° around the axis defined by the D2-D3 arm (Fig. S2c), resulting in the widening angle of about 20° between the two boomerang's arms. The secondary structure of D1 is however, conserved between FljB and FliC (Fig. S2b). Similarly to FliC, it is composed by a long coiled coil flanked by another helix and a β-hairpin which, together with the D0 domain, form the backbone of the flagellar filament. The sequence identity for this domain is 97 % and the Cα traces superpose well with a low r.m.s.d of 0.63 Å for residues 58-177 and 414-458. In domain D2, a short α-helix is formed by FljB residues 309-314, which is not present in FliC. The lower sequence identity for the D2 domain (64%) is reflected by the higher r.m.s.d. of 1.49 Å for the superposition of 123 matching residues. The domain D3 has the lowest sequence identity (30%) and also the greatest r.m.s.d of 1.93 Å, calculated on 87 matching residues, although its topology is well conserved between FljB and FliC (Fig. S2b). Residues 244-249 at one end of the FljB molecule form a short $3_{10}$ helix not present in FliC.

### Supplementary Text S2: Swimming motility is not influenced by flagellin methylation

We investigated if methylation of FliC and FljB would affect flagellar assembly and motility in *S. enterica*. The levels of non-methylated flagellin secreted from a Δ*fliB* mutant strain were comparable to secretion of methylated FliC or FljB (Fig. S4a). Immunostaining of flagella from the WT and a Δ*fliB* mutant strain revealed no significant differences in flagella assembly and flagella numbers per cell body (WT = 2.2 ± 1.8; Δ*fliB* = 2.2 ± 1.5) (Fig. S4b). In agreement with earlier reports [8,9], swimming motility of Δ*fliB* mutant strain in semi-solid agar plates was also not affected (Fig. S4c).

### Supplementary Text S3: Potential role of different methylation patterns found in FliC and FljB

The different methylation patterns found in FljB compared to FliC appear not to play an important role in the ability of the bacteria to adhere to MODE-K cells. Further, the surface hydrophobicity of the respective methylated flagellar filaments appears similar. However, FljB-expressing bacteria are less effective in invading epithelial cells, which has been attributed before to a distinct near-surface swimming behavior of FljB-expressing bacteria [10]. The different orientation of the D3 domain in the two flagellin molecules might in turn modify the shape and the physical-chemical and hydrodynamic properties of the flagellar filament bundle and thus explain the observed distinct motility behavior. As mentioned above, FliC and FljB displayed different methylation patterns and no clear consensus sequence was found for the methylation sites. The observed preferences for small amino acids in the vicinity of the methylation site might favor exposure of the methylated lysines to the epithelial cell surface by avoiding potential sterical interference of bulky side chains. The local amino acid sequence alone, however, does not seem to play a crucial role in this process since most of the methylated, conserved lysines were modified in both fully conserved domains (D0 and D1) and domains (D2 and D3) with low sequence conservation (Supplementary Fig. S2d). Therefore, FliB-dependent methylation seems not to be influenced significantly by changes in the flagellin local structure. This observation suggests that a flagellin assemblies are already present in the bacterial cytosol when FljB and FliC are methylated by FliB before secretion, since methylation sites are found only on the outer surfaces of the flagellar filament (Fig. 3a). Interestingly, we found methylated lysines also in *fliB* deletion mutant bacteria, suggesting that additional methylases are active in the post-translational modification of flagellin.

### Supplementary Figures:

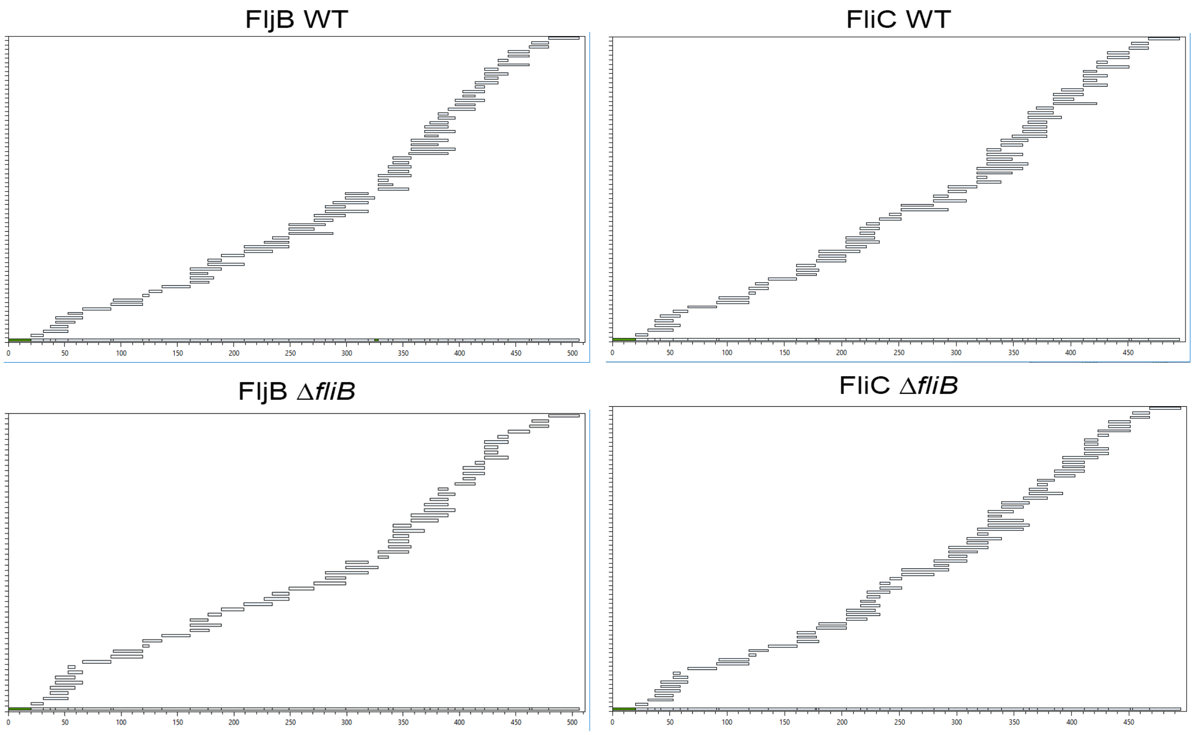


**Supplementary Fig. S1: Sequence coverage maps of mass spectrometry protein methylation studies.** Tryptic peptides that were identified in two of three runs were plotted against the sequence of FljB (left) and FliC (right) for WT (top) and Δ*fliB* samples (bottom). For all proteins the N-terminal sequence (amino acids 1-20) was not covered. Also, amino acids 326-328 in FljB WT are missing. All the other sequence areas are identified, often in multiple peptides, resulting in coverages above 95%. Coverage maps were created using PatternLab software [11].

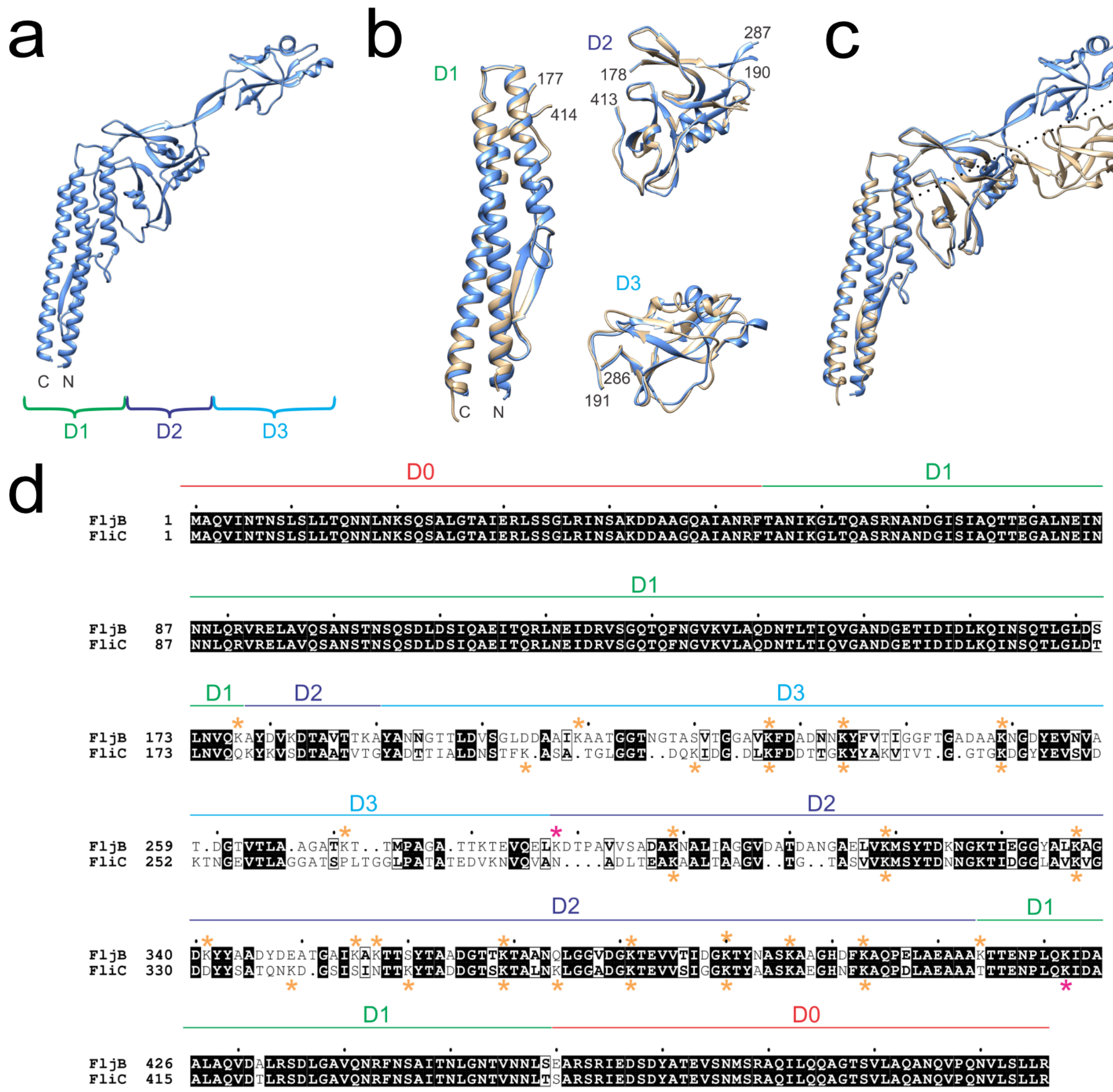


**Supplementary Fig. S2: Structure of truncated FljB, comparison with truncated FliC and methylated lysine residues in the two flagellins of *Salmonella* Typhimurium.** (a) Cartoon representation of truncated FljB. The position of the N- and C-termini is shown at the end of the coiled-coil, and the extension of the domains D1, D2 and D3 is indicated below the structure. (b) Structural superposition of the individual domains. FljB domains are represented in blue, FliC in beige. The N- and C-termini of the structure are indicated, as well as the FljB numbering of the residues at the ends of the polypeptide segments defining the domains. (c) The structures of truncated FljB (blue) and truncated FliC (beige) have been superposed according to the D1 and D2 domains. The domain D3 shows a rotation of about 90° around an ideal axis starting from one end of the coiled-coil in D1 and passing through D2 (shown as dotted line). (d) Alignment of FljB and FliC and methylation pattern. Residues of domains D1, D2 and D3 have been aligned based on the structural superposition obtained with the EBI PDBeFold v2.59 server (51), either superposing the D1-D2 domains or the D3 domain. Residues of domain D0 have been aligned based on the sequence. The extent along the sequence of the domains is indicated above the alignment. Methylated lysine residues are indicated with stars above or below the alignment for FljB or FliC, respectively. Orange stars indicate FliB-dependent methylation sites, magenta stars indicate lysines found methylated both in the WT and in the Δ*fliB* mutant.

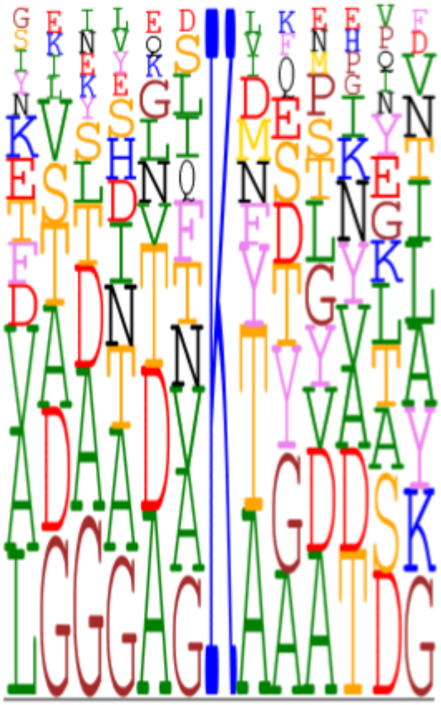

**Supplementary Fig. S3**: **MultiDisp representation of the alignment of methylation sites found in FljB and FliC.** MultiDisp (http://structure.bmc.lu.se/MultiDisp) was used to analyze the conservation of methylation sites (methylated lysine ± 6 residues) in FljB and FliC. The occurrence of the amino acids is proportional to the dimension of the letters.

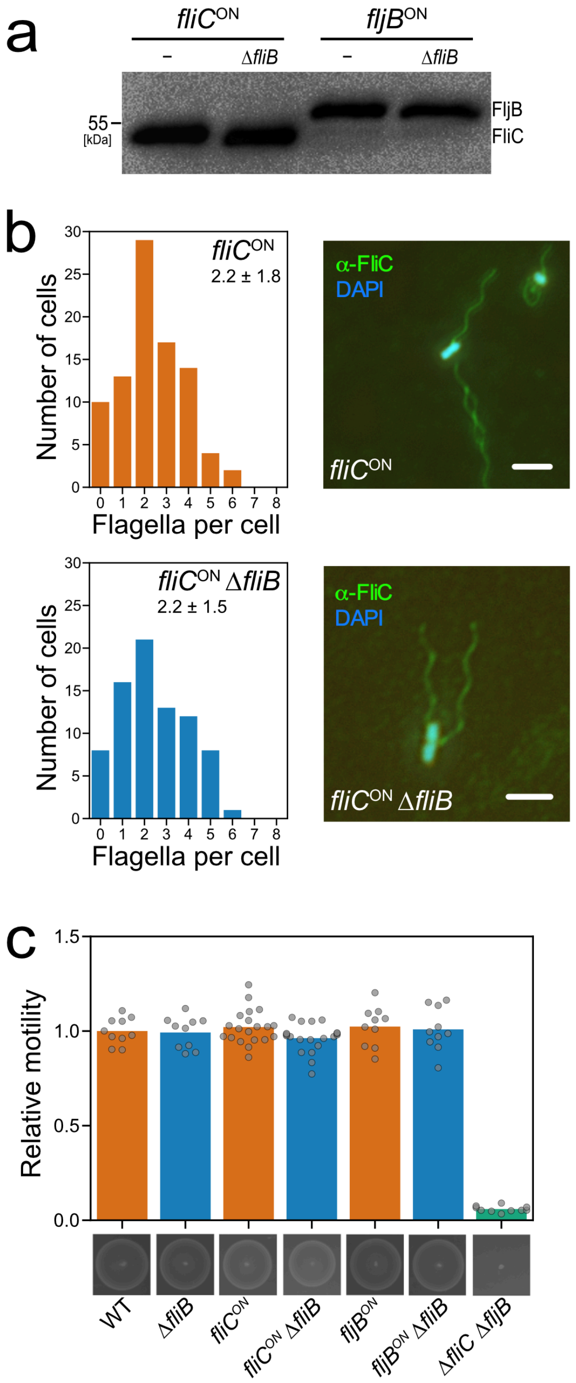


**Supplementary Fig. S4: Swimming motility and flagellar assembly are independent of flagellin methylation.** (a) Immunoblot of secreted flagellins from culture supernatants of *S. enterica*. Secreted proteins were precipitated by addition of 10% TCA and fractionated according to their molecular weight by SDS-PAGE.

Immunoblotting was performed using α-FliC/FljB antibodies. (b) Left: Histograms of the number of flagella per bacterium of FliC-locked strains in the presence or absence of *fliB*. Average flagella numbers were calculated by Gaussian non-linear regression analysis. Right: Flagella filaments were immunostained using α-FliC primary and α-rabbit conjugated AlexaFluor 488 secondary antibodies (green). DNA was stained with DAPI (blue). Scale bar = 5 $\mu$m. (c) Motility phenotypes were analyzed in soft-agar plates containing 0.3% agar and quantified after 4h incubation at 37 °C. Bottom: representative motility plate. Top: The diameters of the motility swarm were measured and normalized to the control strain. The bars display the means of more than 10 biological replicates. Replicates are shown as individual data points.

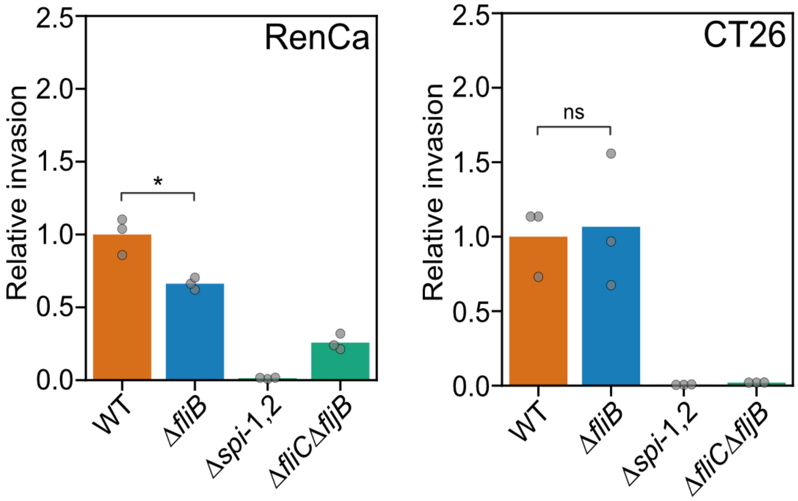


**Supplementary Fig. S5: Relative invasion rates of different eukaryotic host cell types.** The murine epithelial-like cell line RenCa and the murine fibroblast cell line CT26 were infected with *Salmonella* flagella methylation mutants as described for Fig. 2. The bar graphs represent the mean of the reported relative invasion rate data normalized to the inoculum. Replicates are shown as individual data points and statistical significances were determined by the Student's *t* test (* = P<0.05; ns = not significant).

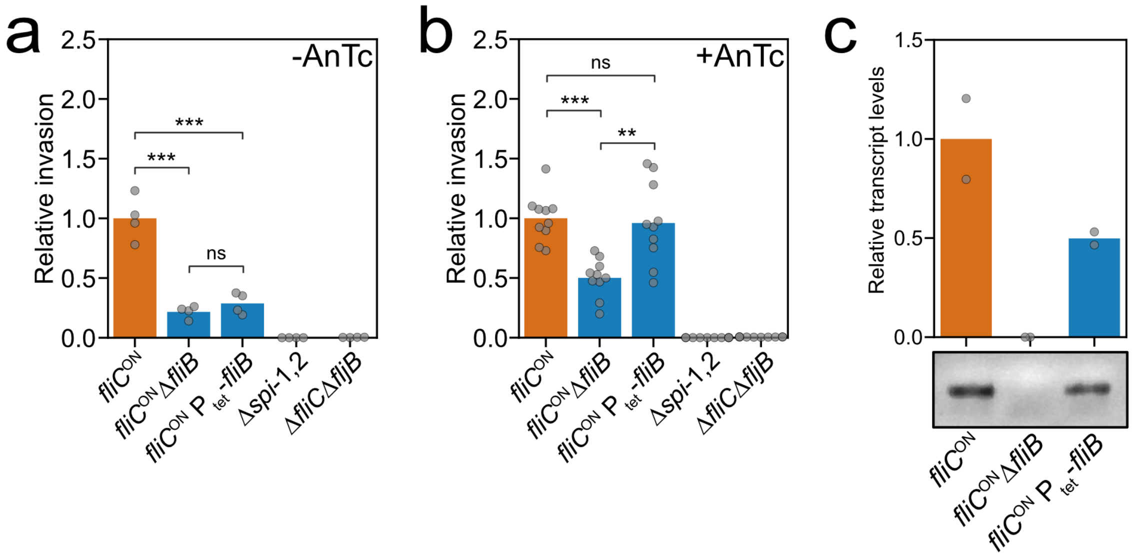


**Supplementary Fig. S6: Complementing the flagella methylation-dependent invasion phenotype by inducible expression of *fliB*.** (a-b) Cell invasion was complemented using an inducible $P_{tet}$ promoter chromosomally fused to *fliB* (EM5080). FliB expression was induced by addition of 100 ng/mL anhydrotetracycline (AnTc). The bar graphs represent the mean of the reported relative invasion rate data normalized to the inoculum. Replicates are shown as individual data points and statistical significances were determined by the Student's *t* test (*** = P<0.001; ** = P<0.01; ns = not significant). (c) Relative *fliB* gene expression of a *fliC*-locked control strain (EM1012), a *fliB* deletion strain (EM4113), and a *fliB*-inducible complementation strain (EM5080) was quantified using qRT-PCR. A representative 1% agarose gel of the amplified *fliB* PCR products is shown below.

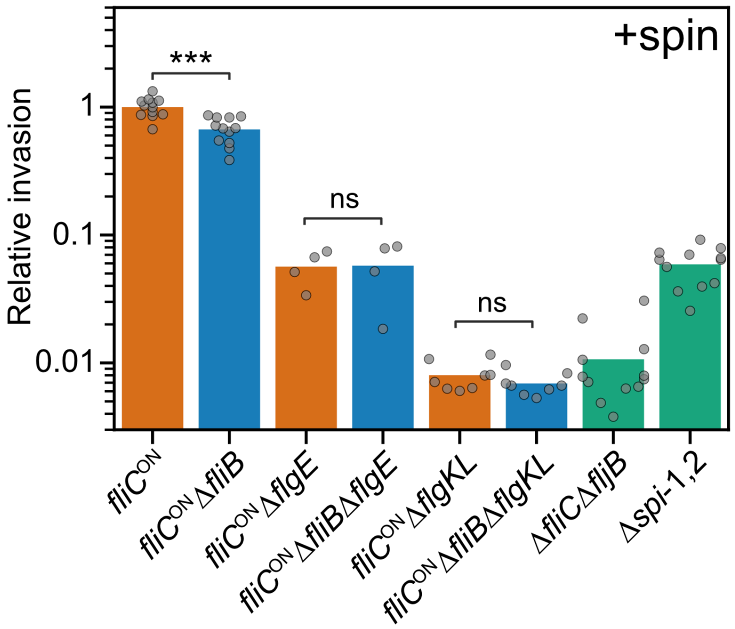


**Supplementary Fig. S7: Flagellin methylation enhances invasion in a flagella-dependent manner.** Infection of MODE-K murine epithelial cells was performed with various flagella assembly mutants at a MOI of 10 for 1h at 37 °C using centrifugation to force contact of the bacteria with the epithelial cells (+spin). Extracellular bacteria were killed by addition of gentamicin for 1 h. Cell lysates were plated in serial dilutions for CFU assessment. Bars represent the mean of the reported relative invasion rate data normalized to the inoculum. Replicates are shown as individual data points and statistical significances were determined by the Student's *t* test (*** = $P<0.001$; ns = not significant).

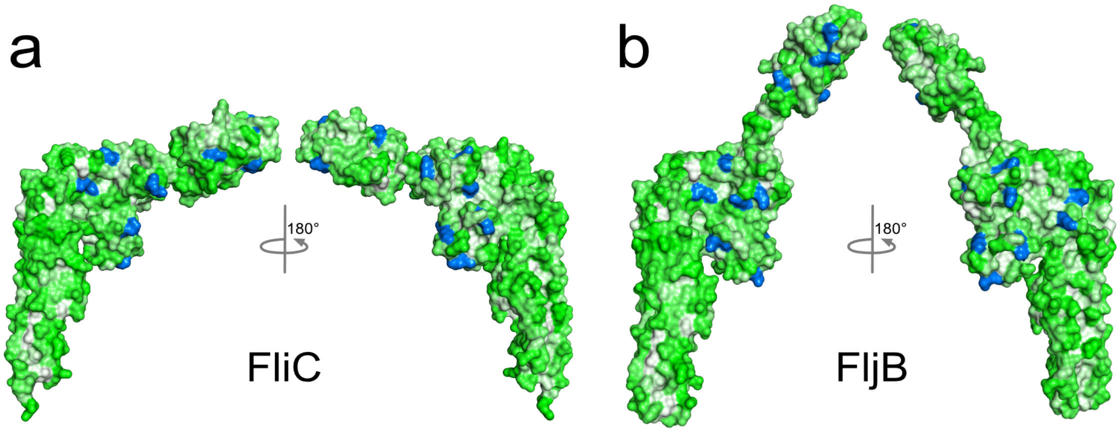


**Supplementary Fig. S8: Surface hydrophobicity distribution of flagellins.** Connolly surface representation of the flagellins FliC (a) and FljB (b). Surface color represents the surface hydrophobicity (green to white indicates increasing hydrophobicity) and methylated lysine residues are highlighted in blue.

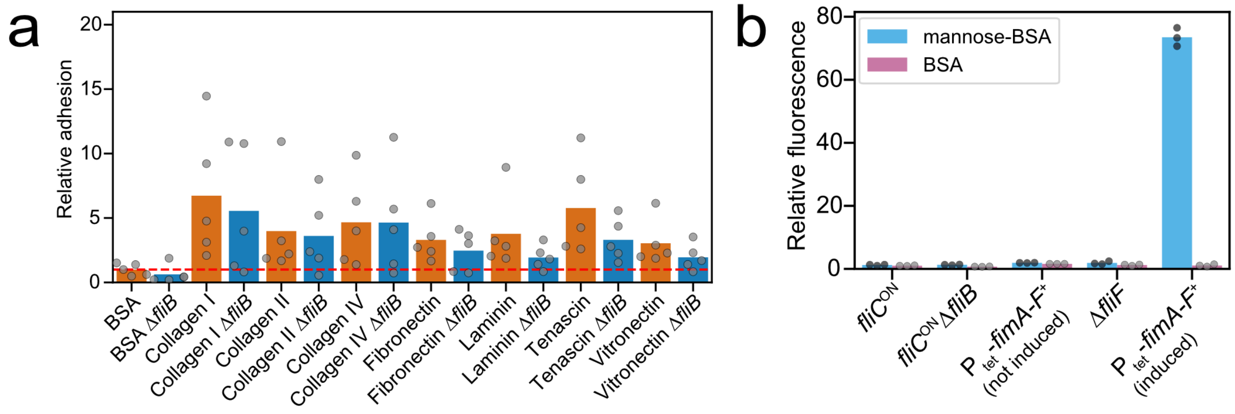


**Supplementary Fig. S9: Flagella methylation does not mediate binding of *Salmonella* to extracellular matrix proteins or mannose residues.** (a) Adhesion of *S. enterica* to extracellular matrix (ECM) proteins *in vitro*. Control and Δ*fliB* strains were added to ECM protein pre-coated 96-well plates and incubated for 1h at 37 °C. After extensive washing, adherent bacteria were plated in serial dilutions for CFU assessment. (b) Methylated flagella do not contribute to binding of *Salmonella* to surface-exposed mannose. Plastic surfaces were coated with BSA or mannose-BSA and strains harboring pFU228-$P_{gapdh}$-mCherry were added for 1 h at 37 °C. After washing, fluorescence was measured in a microplate reader and values were normalized to BSA. $P_{tet}$-*fimA-F* served as positive control after induction with 100 ng/mL anhydrotetracycline. The bar graphs represent the mean of the reported data and replicates are shown as individual data points.

### Supplementary Tables:

**Supplementary Table S1: Data collection and refinement statistics for FljB$_{55\text{-}462}$.** Values in parentheses are for highest-resolution shell. $R_{free}$ is calculated using 5 % of reflections randomly chosen.

| | |
|---|---|
| **Data collection** | |
| Space group | C2 |
| Cell dimensions | |
| a, b, c (Å) | 97.47, 38.28, 124.69 |
| β (°) | 103.54 |
| Resolution (Å) | 50.00 – 2.00 (2.05 – 2.00) |
| $R_{sym}$ or $R_{merge}$ | 0.096 (0.673) |
| *I/σ(I)* | 11.25 (2.26) |
| Completeness (%) | 98.7 (86.0) |
| Redundancy | 5.56 (4.65) |
| **Refinement** | |
| Resolution (Å) | 48.11 – 2.00 |
| No. reflections | 30567 |
| $R_{work}/R_{free}$ | 0.213 / 0.259 |
| No. atoms | |
| Protein | 2938 |
| Water | 299 |
| *B*-factors | |
| Protein | 34.63 |
| Water | 35.98 |
| R.m.s. deviations | |
| Bond lengths (Å) | 0.008 |
| Bond angles (°) | 1.3 |
| Ramachandran plot | |
| Favored | 98.0 % |
| Allowed | 1.5 % |
| Outliers | 0.5 % |

**Supplementary Table S2:** ScanProsite normalized scores of methylation sites matching the Big-1 domain profile. The table includes only the sites matching the Big-1 domain profile. A sequence of 13 residues was scanned for each methylation site. The methylated lysine at the center of each sequence is indicated in bold. Numbering of the sites refers to the position in the sequences reported in Supplementary Fig. S2d. The normalized score reported by ScanProsite measures the confidence level. All the sites fall into the “twilight zone” of confidence (score between 4.1 and 8.5) where true and false positive can co-exist.

| Site | Sequence | Score |
|---|---|---|
| FljB 2 | LDDAAI**K**AATGGT | 4.107 |
| FljB 3 | VTGGAV**K**FDADNN | 4.541 |
| FljB 5 | TGADAA**K**NGDYEV | 4.293 |
| FljB 6 | LAAGAT**K**TTMPAG | 4.396 |
| FljB 7 | TEVQEL**K**DTPAVV | 4.334 |
| FljB 8 | VVSADA**K**NALIAG | 4.603 |
| FljB 9 | NGAELV**K**MSYTDK | 4.128 |
| FljB 13 | TGAIKA**K**TTSYTA | 4.706 |
| FljB 14 | AADGTT**K**TAANQL | 4.892 |
| FljB 15 | LGGVDG**K**TEVVTI | 4.293 |
| FljB 19 | LAEAAA**K**TTENPL | 4.582 |
| FliC 1 | LDNSTF**K**ASATGL | 5.223 |
| FliC 4 | FDDTTG**K**YYAKVT | 5.429 |
| FliC 5 | VTGGTG**K**DGYYEV | 5.140 |
| FliC 6 | ADLTEA**K**AALTAA | 4.933 |
| FliC 7 | GTASVV**K**MSYTDN | 4.231 |
| FliC 9 | YSATQN**K**DGSISI | 4.995 |
| FliC 10 | ISINTT**K**YTADDG | 4.272 |
| FliC 11 | ADDGTS**K**TALNKL | 4.417 |
| FliC 13 | LGGADG**K**TEVVSI | 4.148 |
| FliC 16 | TENPLQ**K**IDAALA | 4.128 |

**Supplementary Table S3:** *Salmonella enterica* serovar Typhimurium SL1344 strains used in this study.

| Strains | Relevant characteristics | Reference/source |
|---|---|---|
| EM774 | SL1344 WT | Lab collection |
| EM812 | SL1344 Δ*fliC7716* Δ*fljB22314* | Lab collection |
| EM1012 | SL1344 Δ*hin-5717*::FRT (*fliC*$^{ON}$) | Lab collection |
| EM1013 | SL1344 Δ*hin-5718*::FRT (*fljB*$^{ON}$) | Lab collection |
| EM1022 | SL1344 Δ*invH-sprB*::FRT (Δ*spi-1*) Δ*sseA-ssaU*::FCF (Δ*spi-2*) | (34) |
| EM1398 | SL1344 Δ*hin-5717*::FRT (*fliC*$^{ON}$) Δ*att*P22::FKF | (34) |
| EM3122 | SL1344 Δ*invH-sprB*::FKF (Δ*spi-1*) | This study |
| EM3123 | SL1344 Δ*hin-5717*::FRT (*fliC*$^{ON}$) Δ*invH-sprB*::FKF (Δ*spi-1*) | (34) |
| EM3124 | SL1344 Δ*hin-5718*::FRT (*fljB*$^{ON}$) Δ*invH-sprB*::FKF (Δ*spi-1*) | (34) |
| EM3734 | SL1344 Δ*fliB8191* | This study |
| EM3758 | SL1344 Δ*fliB8191* Δ*invH-sprB*::FKF (Δ*spi-1*) | This study |
| EM3759 | SL1344 Δ*fliB8191* Δ*hin-5717*::FCF (*fliC*$^{ON}$) Δ*invH-sprB*::FKF (Δ*spi-1*) | This study |
| EM3760 | SL1344 Δ*fliB8191* Δ*hin-5718*::FCF (*fljB*$^{ON}$) Δ*invH-sprB*::FKF (Δ*spi-1*) | This study |
| EM4113 | SL1344 Δ*fliB8191* Δ*hin-5717*::FRT (*fliC*$^{ON}$) | This study |
| EM4114 | SL1344 Δ*fliB8191* Δ*hin-5718*::FRT (*fljB*$^{ON}$) | This study |
| EM4660 | SL1344 Δ*fliB8191* Δ*hin-5717*::FRT (*fliC*$^{ON}$) Δ*att*P22::FCF | This study |
| EM4699 | SL1344 Δ*hin-5717*::FRT (*fliC*$^{ON}$) Δ*flgE6231*(Δaa R106-Q112)::*tetRA* | This study |
| EM4700 | SL1344 Δ*hin-5717*::FRT (*fliC*$^{ON}$) Δ*fliB8191* Δ*flgE6231*(Δaa R106-Q112)::*tetRA* | This study |
| EM4934 | SL1344 Δ*motAB*::*tetRA* | Lab collection |
| EM4949 | SL1344 Δ*hin-5717*::FRT (*fliC*$^{ON}$) Δ*flgKL5739*::FKF | This study |
| EM4950 | SL1344 Δ*hin-5717*::FRT (*fliC*$^{ON}$) Δ*fliB8191* Δ*flgKL5739*::FKF | This study |
| EM5080 | SL1344 *fliB22903*::Tn*10d*Tc (*tetR* terminator) Δ*hin-5717*::FCF (*fliC*$^{ON}$) | This study |

| | | |
|---|---|---|
| EM7748 | SL1344 $P_{tetA}$-*fimAICDHF* / pFU228 ($P_{gapdh}$-*mCherry*) | This study |
| EM7816 | SL1344 Δ*hin-5717*::FRT (*fliC*$^{ON}$) / pFU228 ($P_{gapdh}$-*mCherry*) | This study |
| EM7817 | SL1344 Δ*hin-5717*::FRT (*fliC*$^{ON}$) Δ*fliB8191* / pFU228 ($P_{gapdh}$-*mCherry*) | This study |
| EM7850 | SL1344 Δ*fliF5629*::FKF / pFU228 ($P_{gapdh}$-*mCherry*) | This study |
| EM7842 | SL1344 Δ*fliB8191* Δ*motAB*::*tetRA* | This study |
